# Spatial Transcriptomics-Aided Localization for Single-Cell Transcriptomics with STALocator

**DOI:** 10.1101/2024.06.03.597193

**Authors:** Shang Li, Qunlun Shen, Shihua Zhang

**Author notes:** These authors contributed equally to this work (Shang Li, Qunlun Shen).

## Abstract

Single-cell RNA-sequencing (scRNA-seq) techniques can measure gene expression at the single-cell resolution but lack spatial information. The spatial transcriptomics (ST) techniques simultaneously provide gene expression data and spatial information. However, the data quality on the spatial resolution or gene coverage is still much lower than the single-cell transcriptomics data. To this end, we develop a Spatial Transcriptomics-Aided Locator for single-cell transcriptomics (STALocator) to localize single cells to corresponding ST data. Applications on simulated data showed that STALocator performed better than other localization methods from different angles. When applied to human brain scRNA-seq data and dorsolateral prefrontal cortex 10x Visium data, STALocator could robustly reconstruct the laminar organization of layer-associated cell types. Applications on scRNA-seq data and Spatial Transcriptomics data of human squamous cell carcinoma illustrated that STALocator could robustly reconstruct the relative spatial relationship between tumor-specific keratinocytes, microenvironment-associated cell populations, and immune cells. Moreover, STALocator could enhance gene expression patterns for Slide-seqV2 data and predict genome-wide gene expression data for FISH data, leading to the identification of more spatially variable genes and more biologically relevant GO terms compared to raw data.

## Introduction

Single-cell transcriptomics techniques can measure gene expression at the single-cell resolution, providing deep insights into biological problems such as the development process of organisms and the mechanism of complex diseases. However, single-cell RNA-sequencing (scRNA-seq) data lack spatial information, which is critical for understanding cell differentiation, cancer microenvironment, and brain structure. The newly developed spatial transcriptomics (ST) technologies in recent years can measure gene expression while preserving spatial information, which can provide discovery into some biological problems, such as the regeneration mechanism of planarian^1^, development of the human heart^2^, and analysis of cancer microenvironment^3^.

However, ST technologies still have some limitations. Spatial Transcriptomics^4^ and 10x Visium^5^ data can only be at low resolution, i.e., each spot consists of multiple cells. High-resolution sequencing-based ST data, such as Slide-seqV2^6^, have low data quality. Imaging-based ST data, such as FISH^7^, measure much lower gene numbers than sequencing-based ST data.

Integrating scRNA-seq data and ST data can combine the respective advantages of both data. The current research mainly integrates the two kinds of data from two angles. One is cell-type deconvolution, which uses scRNA-seq data to predict the proportion of each cell type in each spot in the ST data. Researchers have developed many methods, e.g., RCTD^8^ and Cell2location^9^. These methods can only provide a prediction of spatial locations at the cell population level rather than the single-cell level. The other is single-cell localization, which uses ST data to predict the spatial locations of each cell in scRNA-seq data. Researchers have developed several methods, e.g., CellTrek^10^ and scSpace^11^. CellTrek first obtains the low-dimensional representation of scRNA-seq data and ST data through Seurat^12, 13^ and uses the low-dimensional representation and spatial location of ST data to train a random forest classifier to predict the spatial location of scRNA-seq data. However, it needs an interpolation process on ST data to generate new spots around the raw spots. Therefore, it can only localize cells to raw and generated spots, resulting in some cells being localized too densely around raw spots. scSpace uses transfer component analysis (TCA)^14^ to obtain the low-dimensional representation of scRNA-seq data and ST data and trains deep neural networks to fit low-dimensional representations and spatial locations of ST data, thereby predicting the spatial locations of scRNA-seq data. However, some studies of deep transfer learning methods show that the performance of TCA on some domain adaptation tasks is not robust^15–17^, resulting in unreasonable localization. Therefore, an accurate and robust single-cell localization method is still urgently needed.

To this end, we develop a deep learning-based spatial localization method STALocator to localize scRNA-seq data to corresponding ST data. In the context of low-resolution ST data such as Spatial Transcriptomics and 10x Visium, STALocator could directly predict spatial location for cells of scRNA-seq data instead of only providing the proportion of cell types for spots of ST data or only mapping cells of scRNA-seq data to spots of ST data. Given the resolution of ST data such as Slide-seqV2 data and FISH data has approached or reached the single-cell level, STALocator could probabilistically map cells of scRNA-seq data to beads/cells of ST data. Specifically, we applied STALocator to simulated and biological datasets such as the human cerebral cortex and squamous cell carcinoma to accurately identify the relative spatial relationship of cell populations. On high-resolution data such as Slide-seqV2 and FISH, the enhanced data also exhibits an increased capacity for identifying spatially variable genes of greater biological significance than raw data.

## Results

### Overview of STALocator

STALocator is a method that spatially localizes scRNA-seq data through integration with ST data. STALocator is a deep learning-based tool consisting of an integration network that integrates scRNA-seq data with ST data and a localization network that predicts spatial location for scRNA-seq data (**Fig. 1a**). Among them, the integration network adopts the modified domain translation networks^18^ equipped with the sliced Wasserstein distance^19^ to align scRNA-seq and ST data robustly. The localization network adopts the supervised auto-encoder^20^ to fit low-dimensional representations and spatial locations for ST data.

**Figure 1.**
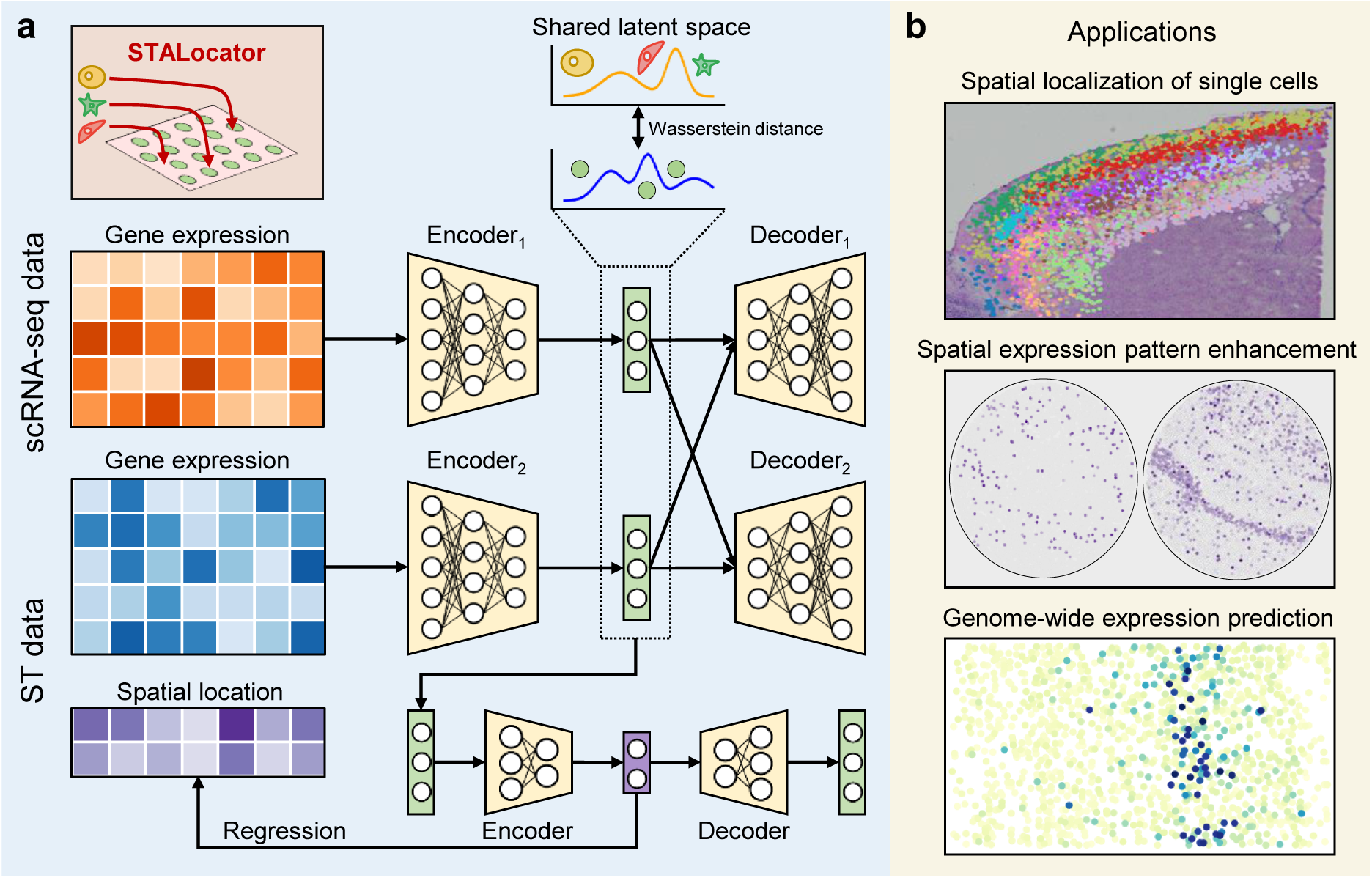
Overview of STALocator. **a,** STALocator takes the top PCs of normalized gene expression profiles of scRNA-seq and ST data as the input of the integration network, obtains batch-free low-dimensional representations in the shared latent space, and fits the spatial locations and low-dimensional representations in the latent space of the localization network. **b,** Applications of STALocator. STALocator can localize the cells of scRNA-seq data to the low-resolution ST data of tissue sections, enhance gene patterns of high-resolution sequencing-based ST data, and predict genome-wide gene expression profiles of imaging-based ST data.

STALocator adopts different strategies for ST data with different resolutions and thus can be applied to different biological scenarios. For low-resolution ST data, such as Spatial Transcriptomics and 10x Visium, STALocator first trains an integration network to obtain a low-dimensional representation that removes batch effects and then trains a localization network to predict the spatial location of cells (**Fig. 1b**). Single-cell data with spatial location information can be considered as a higher-resolution form of spatial transcriptomics data compared to raw ST data. For high-resolution ST data such as Slide-seqV2 and FISH with a resolution closing to scRNA-seq data, STALocator only trains the integration network, then updates the optimal transport (OT) plan during the training process, and finally obtains a global OT plan from scRNA-seq data to ST data which can be used to enhance ST data (**Materials and Methods**). For the Slide-seqV2 data, STALocator enables the acquisition of genome-wide ST data, addressing the limitation of a limited number of measurable genes (**Fig. 1b**).

### STALocator performed better than competing methods in simulation experiments

Considering that the mouse visual cortex STARmap dataset^21^ has a single-cell resolution, we binned them at a specific resolution (900 pixels) to simulate the ST dataset. Taking the raw locations of the raw dataset as ground truth, we localized the raw dataset on the simulated ST dataset by STALocator, CellTrek, and scSpace. STALocator accurately recovered the spatial locations compared with CellTrek and scSpace (**Fig. 2a**). We first observed that STALocator had the lowest mean square error (MSE) on all cells and all seven domains, indicating its superior localization accuracy at the cellular level (**Fig. 2b and Supplementary** Fig. 1b). Furthermore, STALocator had the highest adjusted rand index (ARI), indicating that STALocator more accurately localized the cells of the seven domains to the corresponding domains (**Fig. 2c, d**). Moreover, STALocator had the lowest Kullback-Leibler (KL) divergence on all cells for six of the seven domains, indicating that STALocator outperformed other methods in recovering the actual spatial density distribution of cells (**Fig. 2e, f, and Supplementary** Fig. 1b, c). Finally, STALocator had higher coordinate correlations (**Materials and Methods**) than other methods, indicating that STALocator had the best performance on recovering the marginal distribution of the spatial density of cells along the x-axis and y-axis (**Fig. 2g, h**). Also, we fitted the trends of marker genes from seven distinct domains along the x-axis, confirming the capability of STALocator to accurately retrieve the gene expression patterns associated with these layer-associated marker genes (**Fig. 2i**). In addition, we utilized ARI to test the impact of the sliced Wasserstein distance in the integration network and the decoder in the localization network. Sensitivity and ablation experiments showed these components contributed to the localization performance (**Supplementary** Fig. 1d, e).

**Figure 2.**
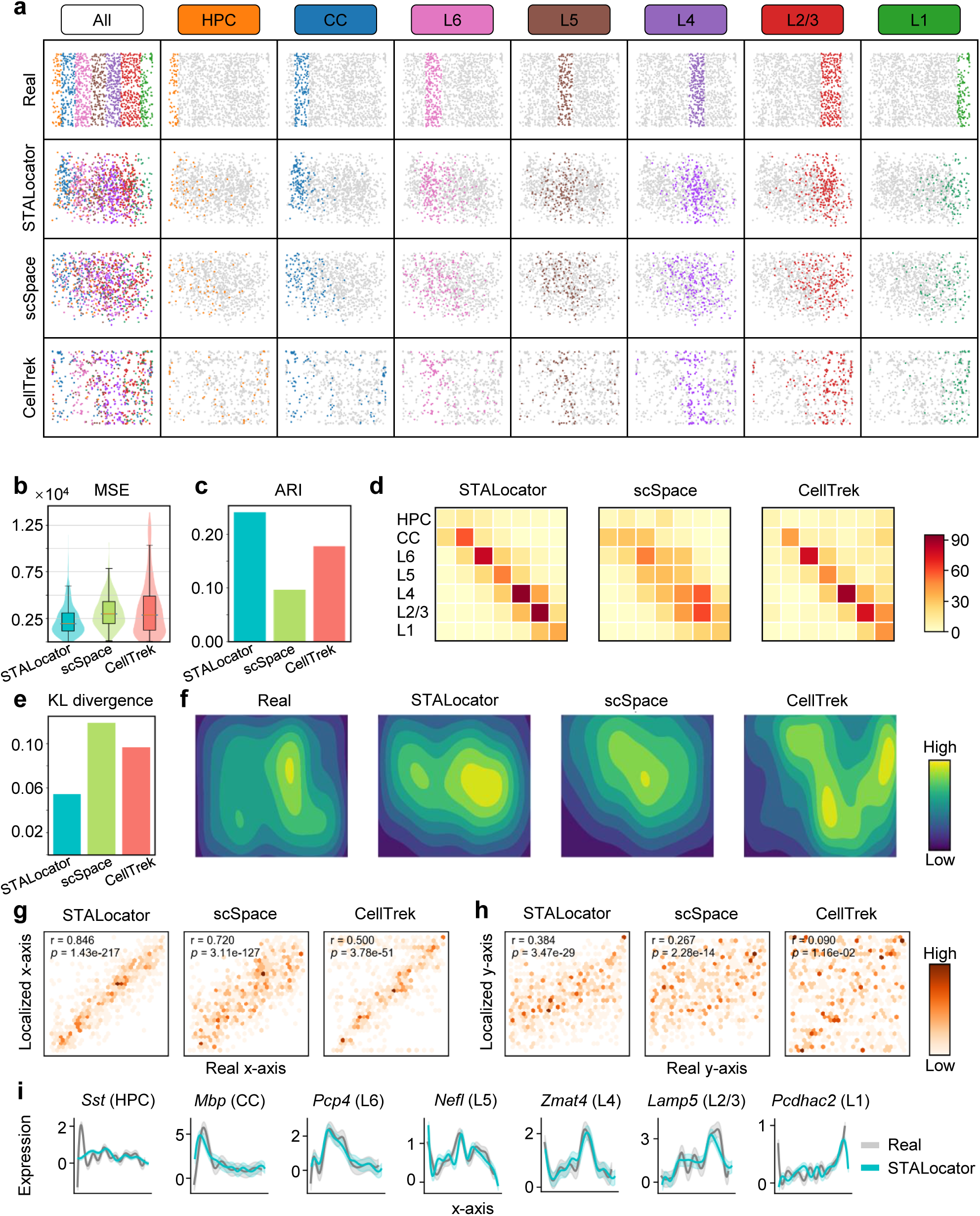
STALocator can accurately recover the spatial locations of the mouse visual cortex STARmap dataset on the simulation experiments. **a,** Ground-truth annotation of seven domains and localization results for different methods. **b,** Boxplot and density curve of localization error in terms of mean square error (MSE) for different methods. **c,** Bar chart of localization accuracy measured by adjusted rand index (ARI) for different methods. **d,** Confusion matrix showing the consistency levels between the localization and raw ones for different methods. **e,** Bar chart of Kullback-Leibler (KL) divergence calculated from spatial grid density for different methods. **f,** Contour plot showing spatial grid density for ground truth and different methods. **g,** Density plot of the correlation test of the x-axis between different methods and ground truth. **h,** Density plot of the correlation test of the y-axis between different methods and ground truth. **i,** The fitted expression trend along the x-axis of marker genes for STALocator and ground truth.

We also applied STALocator to the mouse visual cortex STARmap dataset^21^ using a simulated ST dataset at a higher resolution (700 pixels). STALocator still accurately reconstructed the cells of seven domains (**Supplementary** Fig. 2a) and performed the best compared to the competing methods (**Supplementary** Fig. 2b-h). STALocator also successfully reconstructed the gene expression patterns of marker genes from seven different domains, as evidenced by the expression trends fitted along the x-axis (**Supplementary** Fig. 2i). We further applied STALocator to the mouse brain cortex seqFISH+ dataset^22^ using a simulated ST dataset at a specific resolution (500 pixels). STALocator consistently outperformed the competing methods (**Supplementary** Fig. 3a-e). All these results demonstrated that STALocator showed distinct generalization performance for different data types in simulation experiments.

### STALocator reconstructed the laminar organization of the human and mouse brain cortex scRNA-seq datasets

To test whether STALocator can reconstruct the laminar organization of the layer-associated cell populations, we applied STALocator to two human brain scRNA-seq datasets^23^ from the middle temporal gyrus (MTG) and primary motor cortex (M1) and the human dorsolateral prefrontal cortex (DLPFC) 10x Visium ST dataset^5^ with 12 sections. We first localized the single cells from M1 (**Fig. 3a-c**) and MTG (**Fig. 3d-f**) into the ST dataset section 151673 with the annotated white matter (WM) and six cortical layers. We compared the localization results of STALocator, CellTrek, and scSpace, alongside the deconvolution result of RCTD as a reference (**Fig. 3a, d**). scSpace only restored the hierarchical relationship between cell populations rather than recovered the outlines of the tissue domains, and CellTrek localized many cells too densely around spots of the tissue section. By contrast, STALocator more accurately localized the cell types corresponding to different layers and more evenly localized the cells to the tissue section. The layer-associated cell populations were mainly localized in the cortical layers, and oligodendrocytes were primarily localized in WM (**Supplementary** Fig. 4a, d). Using the manual annotations as ground truth, STALocator achieved the highest ARI, signifying its superior performance in reconstructing laminar organization (**Fig. 3b, e**). We then applied STAGATE^24^ (a spatial clustering method) to the localized scRNA-seq datasets and summarized the distance between localized cell populations and WM. The UMAP visualization of the spatial-aware low-dimensional representation and the distance between domain-associated cell populations and WM illustrated the sequential laminar organization between Layer 2/3 cell populations, Layer 5/6 cell populations, and oligodendrocytes, one of the major cell types on WM^25^ (**Fig. 3c, f**). In contrast, CellTrek incorrectly reconstructed the relative spatial relationships of some cell populations (such as L4 IT), and scSpace failed to localize oligodendrocytes to the tissue section (**Supplementary** Fig. 4c, f).

**Figure 3.**
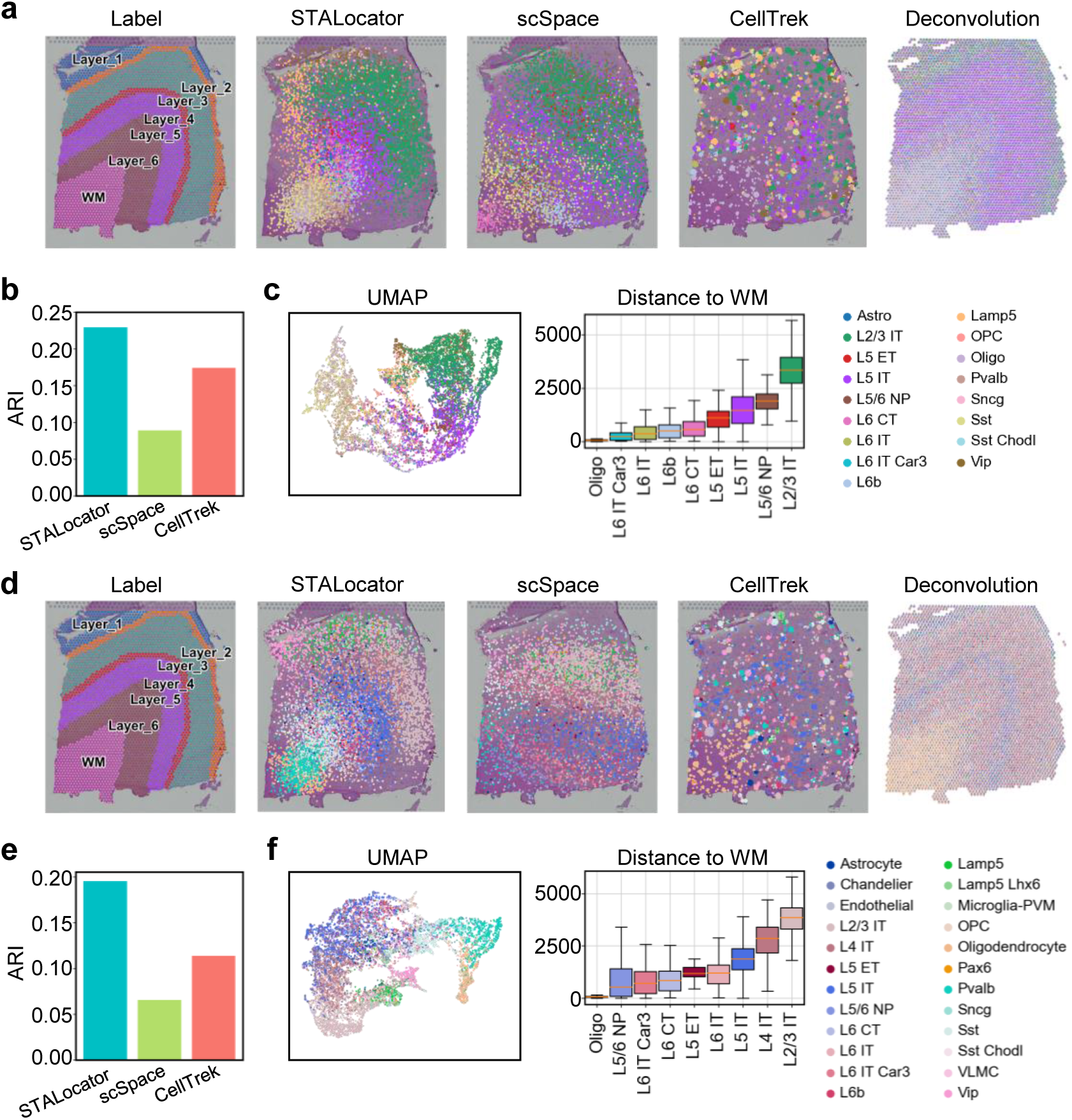
STALocator can better reconstruct the laminar organization of human brain cortex on the human DLPFC dataset (section 151673) with the human brain M1 (a-c) and MTG (d-f) scRNA-seq datasets as paired single-cell data, respectively. **a, d,** Spatial visualization of manual annotation of section 151673, localization results of different methods, and deconvolution results of RCTD. **b, e,** Bar chart of the recovery degree of the laminar organization based on ARI for different methods. **c, f,** UMAP visualizations of the STAGATE-derived low-dimensional representations and boxplot of the distance between WM and selected cell populations of localized scRNA-seq datasets.

When applied to other sections of the DLPFC dataset, STALocator consistently demonstrated superior performance on seven of the 11 sections for the M1 (**Supplementary** Fig. 4b) and MTG (**Supplementary** Fig. 4e) scRNA-seq datasets. We also applied STALocator to the mouse visual cortex scRNA-seq data^26^ and the mouse brain 10x Visium ST dataset sagittal-anterior section 1 (with a part associated with the cortex). STALocator outperformed the two competing methods for reconstructing the hierarchical structure (**Supplementary** Fig. 5a, b). In addition, STAGATE-derived representations of the localized scRNA-seq dataset revealed ordinal laminar organization between Layer 2/3 cell populations and Layer 5/6 cell populations (**Supplementary** Fig. 5c). These results convincingly demonstrated that STALocator can be adapted to different species.

### STALocator reconstructed the relative spatial relationships of cell populations on the human squamous cell carcinoma dataset

The cancer microenvironment contains complex cell-type components and is often closely related to the distance from the cancer focus. We applied STALocator to the paired scRNA-seq and ST datasets collected from the human squamous cell carcinoma (SCC)^3^ tissue and illustrated the histology image and the localization result of STALocator with the cell-type deconvolution result of RCTD as reference (**Fig. 4a**). The results of STALocator suggested that tumor-specific keratinocytes (TSKs) were localized at the leading edge of the cancer region, which was consistent with the original analysis^3^ (**Fig. 4b**). Tertiary lymphoid structure (TLS) consisting of T cells and B cells were primarily localized at tissue domains relatively far from cancer regions than TSKs, which was similar with the cell-type deconvolution result (**Fig. 4c**). Besides, exhausted T cells were predominantly localized at the tissue domains relatively close from cancer regions than naive T cells, which was consistent with the spatial distribution of exhausted scores in the ST data (**Fig. 4d**). These results demonstrated the localization capacity of STALocator.

**Figure 4.**
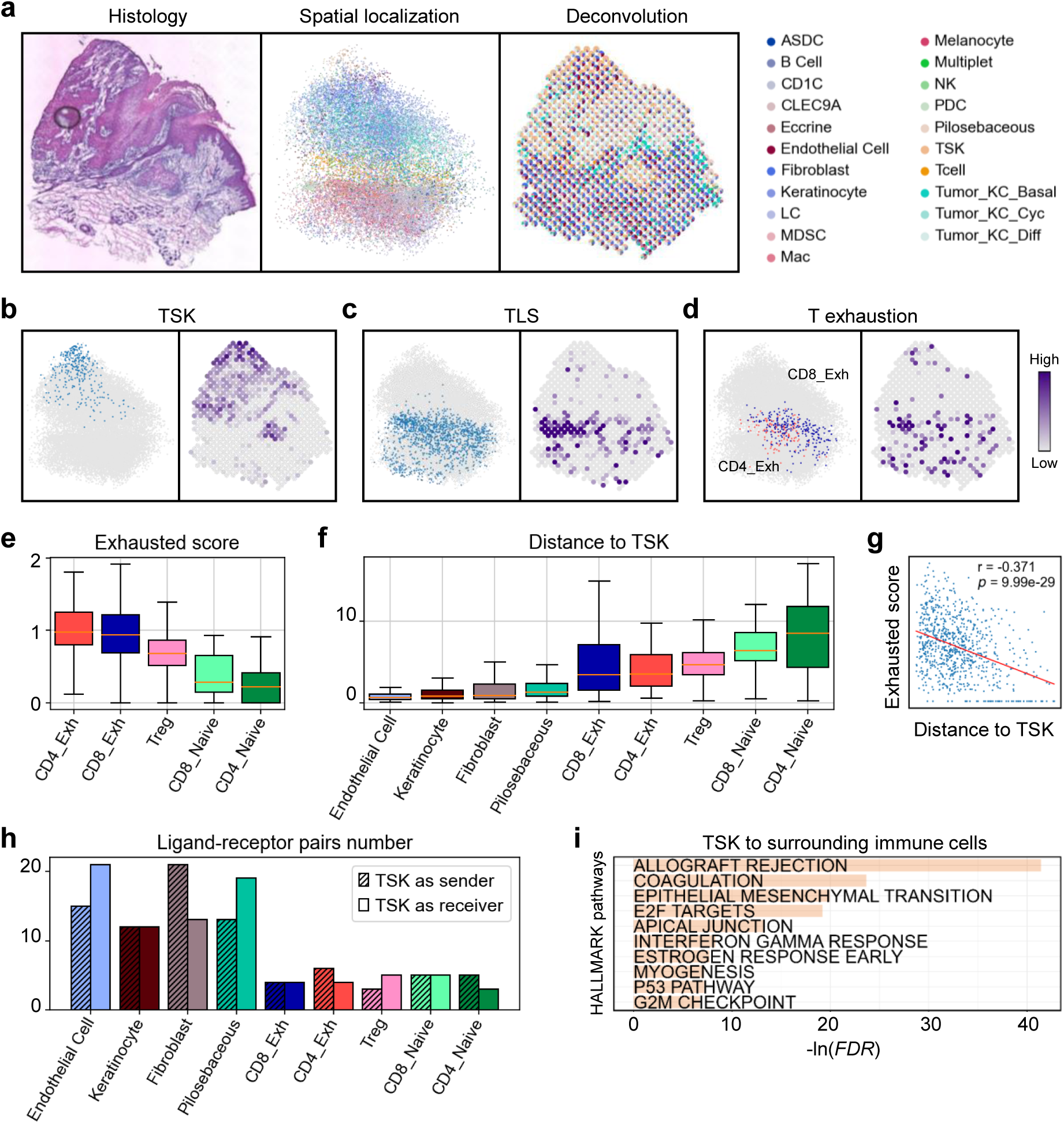
STALocator can robustly reveal the relative spatial relationship between cell populations on the human SCC datasets. **a,** H&E staining image, localization and deconvolution result. **b,** Localization and deconvolution results of Tumor-Specific Keratinocytes (TSKs). **c,** Localization and deconvolution results of Tertiary Lymphoid Structure (TLS). **d,** Localization results of exhausted T cells and exhausted scores of the ST dataset. **e,** Boxplot of exhausted scores of T cell subpopulations. **f,** Boxplot of distance between TSKs and selected cell subpopulations. **g,** Scatter plot showing the relationship of distance to TSK versus exhausted scores of T cell subpopulations. **h,** Number of ligand-receptor pairs between TSK and cell subpopulations selected in **(f)**. **i,** Top ten terms of GSEA result of differentially expressed genes between TSKs and surrounding immune cells.

Compared with naive T cells, exhausted and regulatory T cells had higher exhausted scores and were localized relatively close to TSK (**Fig. 4e, f; Materials and Methods**). That was consistent with the observation that exhausted and regulatory T cells are often present near the tumor leading edges to evade the attack of the immune system^27, 28^. In addition, cell populations such as endothelial cells and fibroblasts were localized relatively closer to TSKs than T cells (**Fig. 4f and Supplementary** Fig. 6a). There was significant negative correlation (r = -0.371, *p* = 9.99e-29) between exhausted scores and spatial distance with TSKs for T cells (**Fig. 4g**). Compared to T cells, cell populations such as endothelial cells and fibroblasts were identified more ligand-receptor pairs, which is consistent with our analysis of the distance between each cell population and TSKs (**Fig. 4h and Supplementary** Fig. 6b). Gene set enrichment analysis (GSEA) of differential expression genes between TSK and surrounding immune cells uncovered many pathways related to immune response on the leading edge of the cancer region, such as interferon-gamma response^29^ (**Fig. 4i; Materials and Methods**). These convincingly demonstrated the capability of STALocator for spatially dissecting the cell populations of the cancer microenvironment.

### STALocator enhanced gene patterns on the mouse brain Slide-seqV2 datasets

Slide-seqV2 data has high spatial resolution but has lower data quality than scRNA-seq data. We applied STALocator to the paired scRNA-seq dataset^30^ and Slide-seqV2 dataset^6^ collected from mouse hippocampus. STALocator could transfer the cell-type labels from the scRNA-seq data to the ST data (**Materials and Methods**) relating to critical domains of the mouse hippocampus supported by the Allen Brain Atlas^31^ (**Fig. 5a, b**). For instance, the CA1, CA2, and CA3 principal cells were primarily distributed within the cornu ammonis (CA) domain, while the dentate gyrus (DG) domain was mainly composed of the dentate principal cells. Based on the enhanced data, marker genes of these structures illustrated clear expression patterns that were consistent with spatial localization of corresponding cell populations (**Fig. 5c**). For example, the expression of *Itpka*, *Amigo2*, and *Hs3st4*, classic marker genes of CA1, CA2, and CA3 subdomains respectively^32–34^ was more distinct than the raw data. Moreover, the enhanced expression pattern of *Calb1*, a known marker gene of the DG domain^35^, was notably clear on the DG domain (**Fig. 5c**). These results demonstrated the ability of STALocator on ST data enhancement.

**Figure 5.**
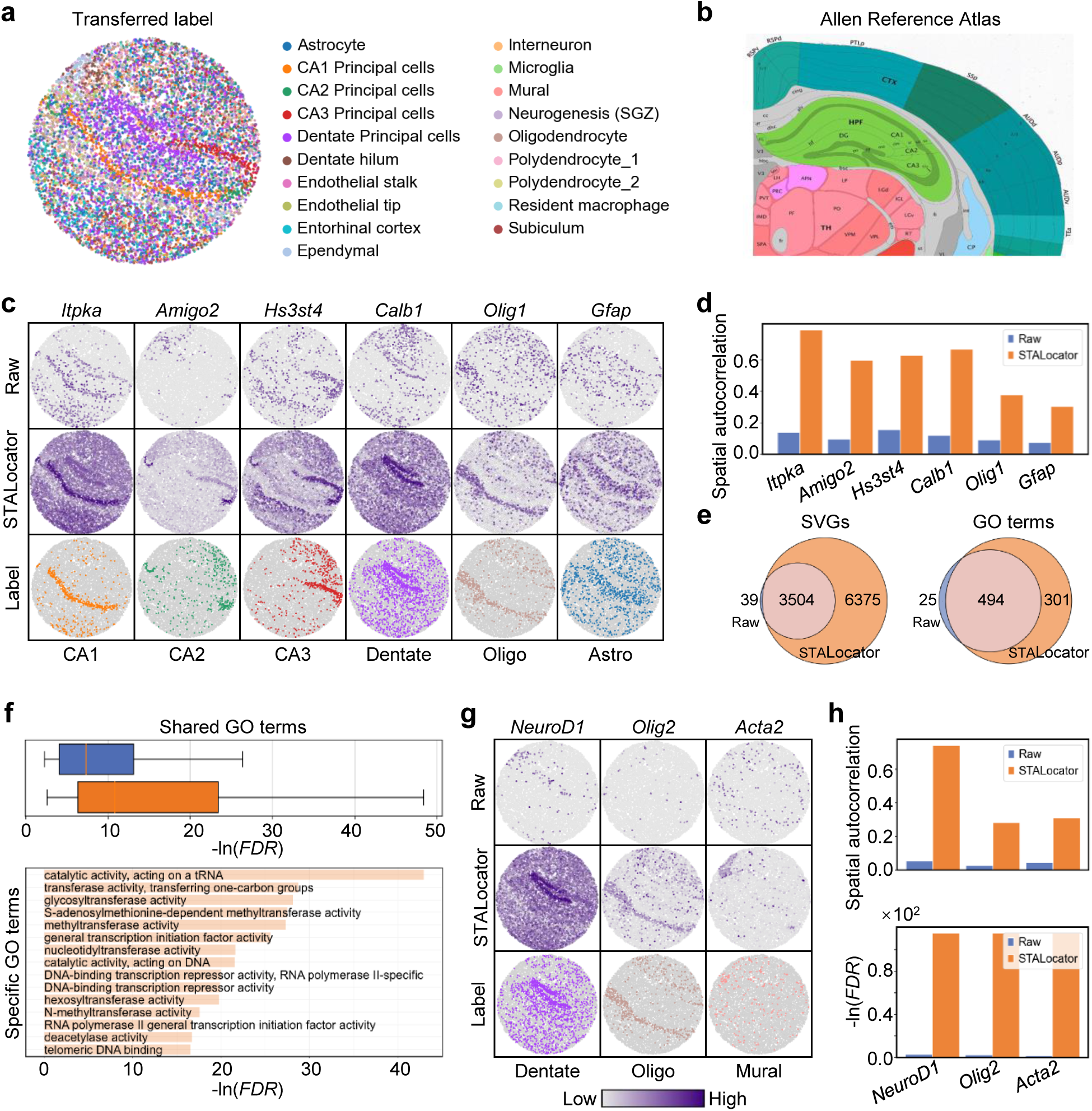
STALocator can enhance gene expression patterns on the mouse hippocampus Slide-seqV2 dataset. **a,** Spatial visualization of cell-type label transferring results. **b,** ISH images from the Allen Brain Atlas. **c,** Spatial visualization of the marker genes for the raw and enhanced dataset and corresponding transferred labels. **d,** Bar chart showing spatial autocorrelations of genes in (**c**) of the raw and enhanced datasets. **e,** Venn plot of the identified SVGs and enriched Gene Ontology (GO) terms of the raw and enhanced data. **f,** Boxplot of the significance of shared GO terms, and bar chart of the top 15 specific GO terms of SVGs from the enhanced data by STALocator. **g,** Spatial visualization of the specific SVGs for the raw and enhanced data by STALocator and corresponding transferred labels, respectively. **h,** Bar chart showing spatial dependency test results of genes in (**g**) of the raw and enhanced data by STALocator (i.e., spatial autocorrelation and FDR).

We identified more SVGs by Hotspot^36^ from the enhanced data than the raw one and observed higher spatial autocorrelation (**Fig. 5d, e**). Moreover, most of the SVGs identified from the raw data were covered by those from the enhanced one, indicating the enhancement by STALocator could help to depict spatial expression specificity (**Fig. 5e**). Enrichment analysis of these two SVG sets showed more significant biological relevance for SVGs from the enhanced data (**Fig. 5f**). Among the shared GO terms, some (e.g., GTPase binding^37^) are related to signal transduction on the brain (**Supplementary** Fig. 7). And SVGs of enhanced data were enriched in several unique GO terms (e.g., telomeric DNA binding^38^) related to neurogenesis (**Fig. 5f**). Many marker genes were only identified as SVGs in the enhanced data and had higher spatial autocorrelations and significance than the raw one (**Fig. 5g, h**). For instance, the enhanced expression pattern of *Olig2*, a known marker gene of oligodendrocytes^39^, reflected the WM domain distinctly (**Fig. 5g**). In short, the enhanced data by STALocator could yield more biological relevance and suggest that STALocator could significantly enhance the data quality of the Slide-seqV2 ST data.

We also applied STALocator to the paired scRNA-seq^40^ and Slide-seqV2 data^6^ from the mouse olfactory bulb tissue. The label transfer coupled with STALocator explicitly revealed the hierarchical structure of the olfactory bulb (**Supplementary** Fig. 8a, b). Enhanced data were identified with more SVGs, which were enriched in more biological meaningful terms, including the majority of SVGs and terms from the raw one (**Supplementary** Fig. 8c). The enrichment analysis of SVGs from the enhanced data also discovered some informative GO terms (e.g., peptidase activator activity^41^) relating to protein metabolism during odor perception (**Supplementary** Fig. 8d). Several marker genes (e.g., *Syt1*, *Sox10*, and *Rbp1*) illustrated more distinct expression patterns in the enhanced data by STALocator (**Supplementary** Fig. 8e).

### STALocator predicted genome-wide gene expression profiles on the mouse brain cortex datasets profiled by FISH

FISH data is another type of high-resolution ST data, notably distinguished by its single-cell resolution. However, because of its technical difficulty and high cost, the number of genes measured is usually much smaller than the sequencing-based data. Here, we applied STALocator to the mouse primary visual area (VISp) scRNA-seq dataset^42^ and mouse visual cortex STARmap dataset^21^ for predicting genome-wide gene expression and revealed several layer-related cell types that were consistent with the original labels through the label transfer procedure (**Fig. 6a, b**). The gene throughput of the raw data was increased from 1,020 to 38,048. From the extended data by STALocator, we identified more SVGs and terms than the raw one, where most of the SVGs and terms from the raw one were retained (**Fig. 6c**). From the extended data, we found several specific GO terms related to neurogenesis and cell migration (e.g., GTPase regulator activity^43^) (**Fig. 6d**) and several biologically relevant shared GO terms relating to the signal transduction on brain (e.g., glutamate binding^44^) (**Supplementary** Fig. 9). The extended data preserved the original expression patterns of measured layer-specific marker genes (**Fig. 6e, f**). Moreover, the predicted expression patterns of other unmeasured marker genes can be verified by staining images^31^ (**Fig. 6g, h**).

**Figure 6.**
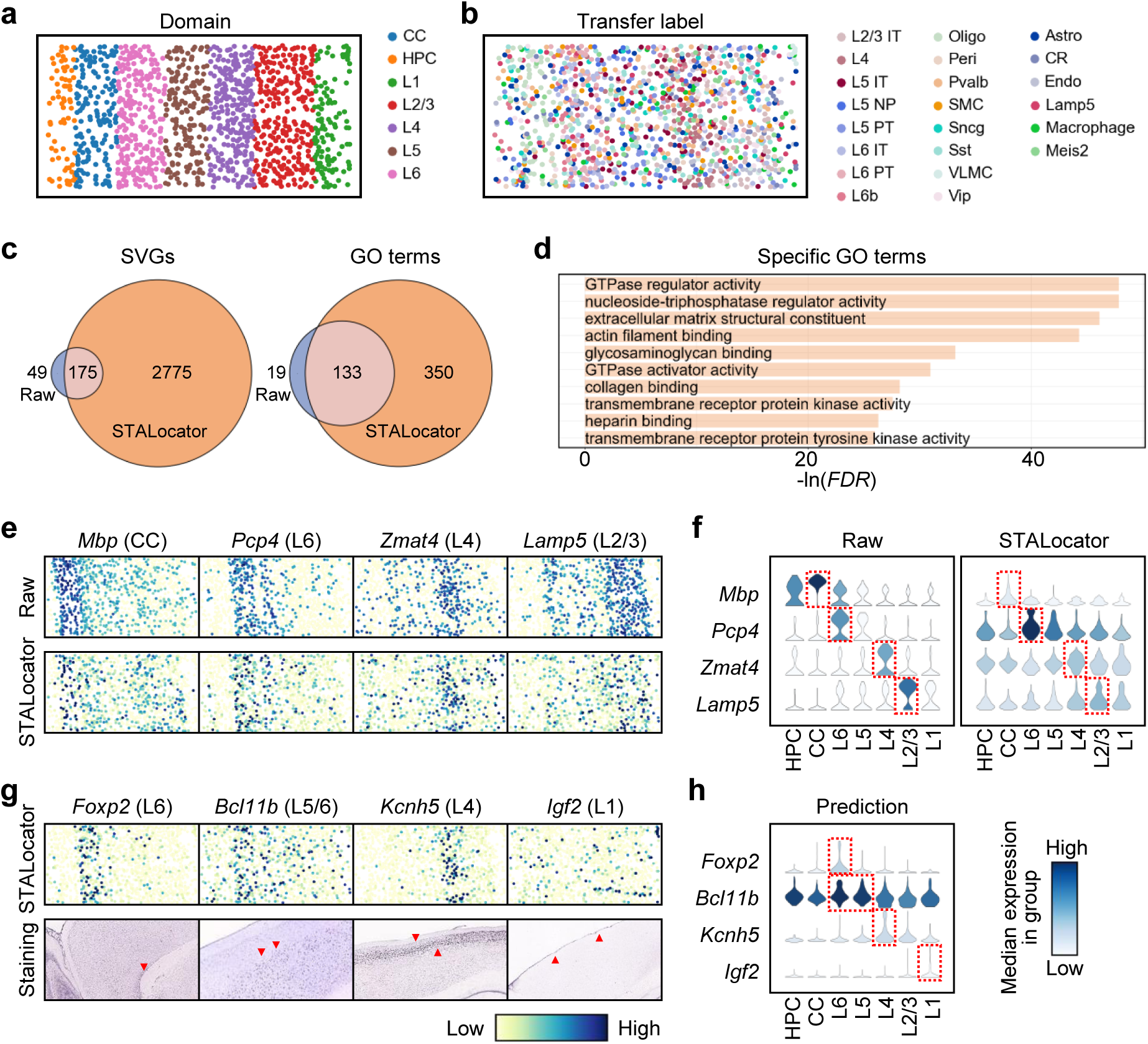
STALocator can predict genome-wide gene expression profiles on the mouse visual cortex STARmap dataset. **a,** Spatial visualization of manual annotation of seven domains. **b,** Spatial visualization of cell-type label transferring results. **c,** Venn plot of the identified SVGs and enriched GO terms of the raw and enhanced data by STALocator. **d,** Bar chart showing the top ten specific GO terms of SVGs from the enhanced data by STALocator. **e,** Raw and recovered expression profiles of the measured marker genes. **f,** Violin plot of the raw and recovered expression of genes in (**e**). **g,** Predicted expression of the unmeasured marker genes and corresponding staining images from the Allen Brain Atlas. **h,** Violin plot of the predicted expression of genes in (**g**).

We further applied STALocator to the mouse VISp scRNA-seq dataset^42^ and mouse brain cortex seqFISH+ dataset^22^ and observed the laminar organization of layer-related cell types by transferring cell-type labels of the single-cell data to the ST data (**Supplementary** Fig. 10a, b). The gene throughput of the raw data was expanded to approximately 3.8 times from 10,000 genes to 38,048 genes by STALocator. We identified more SVGs from the extended data than the raw one with several shared ones, and observed similar results for the enriched GO terms of SVGs (**Supplementary** Fig. 10c). From the extended data, we found several specific GO terms related to the regulation of neuronal activity (e.g., rRNA binding^45^) (**Supplementary** Fig. 10d). For some classic marker genes, the extended data by STALocator retained the original expression patterns of measured genes (**Supplementary** Fig. 10e), and predicted clear expression patterns for many unmeasured genes (**Supplementary** Fig. 10f). Collectively, these results demonstrated the reliability of STALocator in predicting genome-wide expression profiles for FISH data and revealed more biological relevance.

## Discussion

STALocator localizes single cells onto ST data and reveals the relative spatial relationships among cell populations. STALocator compensates for the limitations of ST data in terms of resolution and data quality and imparts spatial information to the scRNA-seq data. That combines the advantages of the two data types and can provide deeper spatial insights into some biological problems.

scSpace failed to remove batch effects robustly, resulting in unreasonable localization. CellTrek is essentially a mapping method that can only map cells to denser spatial coordinates that are augmented in advance, making it not an end-to-end localization method. CellTrek often localizes some cells too densely **(Fig. 3a)**. Moreover, other mapping methods, e.g., Tangram^46^, can only map cells to samples of the ST data, making it hard to predict finer spatial coordinates for scRNA-seq data when applied to the low-resolution ST data. In addition, some deconvolution methods, e.g., Cell2location and RCTD, can only predict cell type fractions of spots of the ST data. In short, STALocator provides a novel tool for integrating scRNA-seq and ST data.

For the high-resolution ST data, considering that the resolution of ST data is close to or has reached the single-cell level, STALocator can map single cells onto the ST data directly. For sequencing-based data, e.g., Slide-seqV2 data, STALocator could enhance the data quality compared to the raw one. For imaging-based data, e.g., FISH data, STALocator can provide genome-wide gene expression profiles. Moreover, the enhanced ST data can help to identify more SVGs with more biological relevance than the raw data.

Since STALocator is a deep learning-based approach, its results often exhibit a certain level of randomness. For low-resolution ST data, STALocator typically takes about 5 minutes to run 10,000 epochs. However, for high-resolution data, due to the incorporation of OT in the training process, STALocator takes over 20 minutes to run 5,000 epochs. Currently, STALocator is limited to the integration between transcriptomics data. However, it is anticipated that STALocator can be expanded to integrate multi-omics data.

## Materials and Methods

### Data description

We applied STALocator to a combination of multiple pairs of scRNA-seq (**Supplementary Table S1**) and ST datasets (**Supplementary Table S2**). The human DLPFC 10x Visium dataset has 12 sections with the spot number ranging from 3,498 to 4,789. We down-sampled the corresponding human brain M1 and MTG scRNA-seq datasets to 10,000 cells each. The human SCC Spatial Transcriptomics dataset has 9 sections from 3 patients, and we selected section 1 from patient 2 with 666 spots. We selected the paired scRNA-seq dataset only from tumor tissue, which included 26,299 cells. We down-sampled the Slide-seqV2 and scRNA-seq datasets of mouse hippocampus to 10,000 beads and 20,000 cells, respectively. The mouse visual cortex STARmap and mouse VISp scRNA-seq dataset included 1,207 and 11,759 cells, respectively.

### Data preprocessing

In both ST and scRNA-seq datasets, we filtered out genes expressed in less than three cells/spots/beads, log-transformed the raw gene expression counts that were normalized by library size, identified highly variable genes (HVGs), scaled the expression matrix by genes, and then performed joint principle component analysis (PCA) using python package SCANPY^47^.

### Model architecture

STALocator consists of two networks: the integration network and the localization network. The two networks are trained separately, with the localization network being trained after the integration network.

#### Integration network

This part integrates the scRNA-seq and ST data (considered as two domains) to obtain low-dimensional representations of cells/spots without batch effects. The integration network adopts two domain translation networks for two batches of data^18^. It takes the top *p* principal components of cells/spots as inputs. Let *M* and *N* be scRNA-seq data and ST data respectively, and *Y* be the two-dimensional physical space of the tissue section of the ST data. Let *X* be the shared latent space of the two domain translation networks. The encoder *E*_1_(·): *M* → *X* can obtain low-dimensional representation *E*_1_(*m*) ∈ *X* for ∀*m* ∈ *M*. And the decoder *D*_2_(·): *X* → *N* can then transfer *E*_1_(*m*) ∈ *X* to *D*_2_(*E*_1_(*m*)) ∈ *N* for ∀*m* ∈ *M*. The encoder *E*_1_(·): *M* → *X* and the decoder *D*_2_(·): *X* → *N* form a domain translation network *D*_2_(*E*_1_(·)): *M* → *N*. Similarly, the encoder *E*_2_(·): *N* → *X* and the decoder *D*_1_(·): *X* → *M* form another domain translation network *D*_1_(*E*_2_(·)): *N* → *M*. Moreover, two discriminators *D*_*m*_(·): *M* → (0,1) and *D*_*n*_(·): *N* → (0,1) distinguish the raw and transferred cells/spots. Let *P*(*E*_1_(*m*)) and *P*(*E*_2_(*n*)) be the empirical distribution of *E*_1_(*m*) and *E*_2_(*n*). The training process is a min-max optimization as follows,

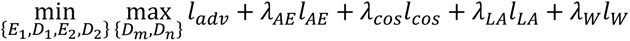

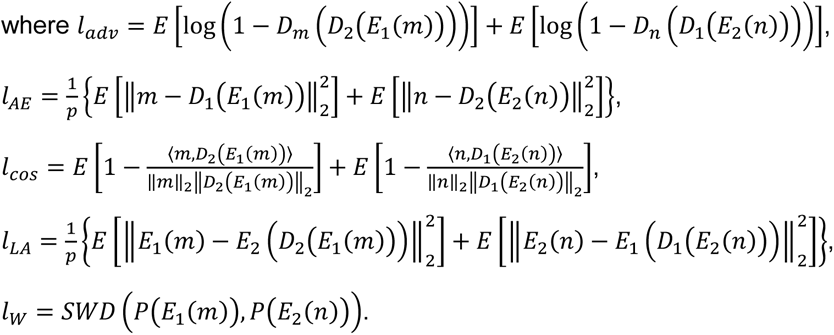

Among these loss terms, *SWD* adopts an approximate Wasserstein distance using the slicing strategy^19^.

For all analyses, we set *λ*_*AE*_ = 10, and *λ*_*LA*_ = 10. For the low-resolution ST data, we set *λ*_*W*_ = 5 and *λ*_cos_ ∈ {2,5,10}. The final choice of *λ*_*cos*_ depends on ARI (when labels are available in ST data) or silhouette coefficient (when labels are not available in ST data). For the high-resolution ST data, we set *λ*_*W*_ = 2, and *λ*_*cos*_ = 15.

#### Localization network

This part predicts the spatial locations of cells. The localization network adopts a spatial location-supervised auto-encoder^20^. The encoder *E*(·): *X* → *Y* takes low-dimensional representations of the integration network as inputs and outputs spatial locations, and the decoder *D*(·): *Y* → *X* reverses them. Only the ST data takes part in the training process, and the trained encoder predicts spatial locations for the scRNA-seq data. Let *S* be the spatial locations of the ST data. For each location s ∈ S, the loss function of the localization network is as follows,

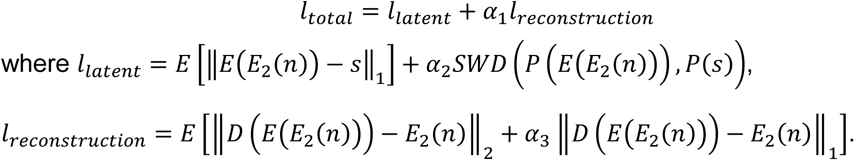

Here, *SWD* is the same as the integration network. For all analyses, we set *α*_1_ = *α*_2_ = *α*_3_ = 0.1.

### Label transfer and data enhancement

For the Slide-seqV2, FISH, and paired scRNA-seq datasets, we performed the minibatch OT^48^ process using the Sinkhorn algorithm^49^ during the training process. Specifically, we computed the cost matrix *C* by a given metric in an epoch using the Python package POT^50^. Let *P* be the OT plan and *B*_*m*_ and *B*_*n*_ be the batch sizes, respectively. We denote the total cost as 〈*C*, *P*〉 ∶= ∑_*i*,*j*_ *C*_*ij*_*P*_*ij*_, the entropy regularization term as *H*(*P*) ∶= − ∑_*i*,*j*_ *P*_*ij*_(log *P*_*ij*_ − 1), and the margin distribution as *a* 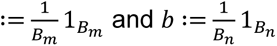, respectively. The optimal OT plan *P*^∗^ can be computed by solving the following optimization problem,

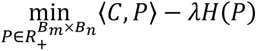

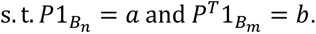

Let denote 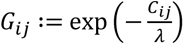 and initialize with 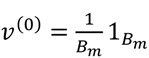. The iteration process is as follows,

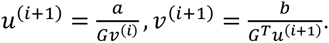

After several iterations, the optimal OT plan can be obtained by *P_ij_*^∗^ = *u*_*i*_*G*_*ij*_*v*_*j*_. We finally obtained the global OT plan to probabilistically map cells of scRNA-seq data to beads/cells of ST data. We averaged the global OT plan by cell types and transferred the labels with maximum probability for ST data. Then, the ST data was enhanced by multiplying normalized scRNA-seq data and the global OT plan.

### ST data deconvolution

For the 10x Visium, Spatial Transcriptomics, and paired scRNA-seq datasets, we predicted cell-type fractions for all spots of the ST data using the R package RCTD^8^ according to its tutorial (https://github.com/dmcable/spacexr). Specifically, we run the *create.RCTD* function with parameters max_cores = 10, UMI_min = 0, UMI_min_sigma = 0, MAX_MULTI_TYPES = number of cell types, CELL_MIN_INSTANCE = 0, and keep_reference = TRUE, and we run the *run.RCTD* function with parameters doublet_mode = ‘full’.

### Spatial trajectory inference

For the human DLPFC and human brain cortex scRNA-seq datasets, we obtained the spatial-aware low-dimensional representations of localized scRNA-seq datasets using the Python package STAGATE^24^ according to its tutorial (https://github.com/zhanglabtools/STAGATE). We performed the PAGA algorithm^51^ to infer spatial trajectory and visualized the PAGA graphs by the *scanpy.pl.paga()* function.

### Calculation of exhausted scores

For the human SCC datasets, let *M*_*a*×*d*_ and *N*_*b*×*d*_ be the normalized gene expression of scRNA-seq and ST dataset, respectively, and *T* be the set of exhaustion-associated genes (i.e., *LAG3*, *TIGIT*, *PDCD1*, *HAVCR2*, *CTLA4*, *BTLA*, *KLRC1*, *ENTPD1*, and *LAYN*) selected from the original study^3^. Exhausted scores were calculated as the average expression of exhaustion-associated genes, that is 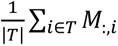 for the scRNA-seq dataset and 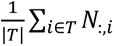 for the ST dataset.

### Intercellular distance analysis

For the human SCC datasets, we calculated the Euclidean distance between cells on the localized scRNA-seq dataset. For each cell population, we defined the median distance between each cell and the nearest TSK as its ‘distance from the TSKs’. We then defined T cells and B cells as immune cells. Immune cells whose distance from the nearest TSK is less than a fixed threshold (we set it as 1, close to the diameter of spots) are defined as ‘surrounding immune cells’.

### Identifying ligand-receptor pairs

For the human SCC datasets, we identified ligand-receptor pairs between cell population pairs using the R package CellChat^52^ following its tutorial. The results included the communication strength and ligand-receptor pairs numbers of cell population pairs.

### Identifying spatially variable genes

For the Slide-seqV2, FISH, and paired scRNA-seq datasets, we identified SVGs using the Python package Hotspot^36^ following its tutorial (https://github.com/Yoseflab/Hotspot). The results included the spatial autocorrelation and *p*-value of each gene. The genes with FDR < 0.001 were considered as SVGs.

### Biological process enrichment analysis

For the human SCC datasets, we first performed differential expression analysis between TSKs and the surrounding immune cells using the *FindMarkers* function and ranked the differentially expressed genes using the *avg_log2FC* function in the Seurat package. We selected the HALLMARK pathway list and performed the GSEA enrichment analysis using the *fgseaMultilevel* function of the R package fgsea^53^. The results included the *p*-value and normalized enrichment score (NES) of each pathway.

For the Slide-seqV2, FISH, and corresponding scRNA-seq datasets, we performed the GO enrichment analysis on identified SVGs using the *enrichGO* function of the R package clusterProfiler^54^. The pathways with *FDR* < 0.05 were considered as significant enrichment.

### Benchmarking localization methods

We compared STALocator with two competing localization methods, i.e., CellTrek and scSpace. We predicted specific spatial locations for cells and filtered out the cells outside the tissue area. The implementation of these methods is as follows.

#### CellTrek

CellTrek is a pseudo-localization method based on the Seurat integration and RF model. We performed CellTrek on the simulation experiments and the human DLPFC 10x Visium dataset with paired scRNA-seq datasets. We ran the model using the *celltrek* function with parameters intp_pnt=5000, dist_thresh=0.55, spot_n=30 (10x Visium data) or maximum number of cells included in pseudo-spots (simulated data), top_spot=1, repel_r=20, repel_iter=20.

#### scSpace

scSpace is an end-to-end localization approach based on TCA and deep neural networks. Following the standard workflow, we performed scSpace on the simulation experiments and the human DLPFC 10x Visium dataset with paired scRNA-seq datasets. We preprocessed the data using the *preprocess* function with parameters n_features=2000 (10x Visium data) or the number of genes (simulated data) and select_hvg=‘union’. We ran the model using the *construct_pseudo_space* function with parameters batch_size=128 (10x Visium data) or 16 (simulated data).

### Evaluation metrics

We designed the following metrics to evaluate localization performance.

#### Mean square error (MSE)

We used MSE for the simulation experiments and calculated it from the Euclidean distance between the localized coordinates and raw coordinates of cells. It can measure the error between localization results and ground truth at the single-cell level.

#### Adjusted rand index (ARI)

For the simulation experiments, we trained a k-NN classifier using the cell-type labels and raw coordinates of the raw data and predicted cell-type labels for the localized coordinates of single-cell data. For the human DLPFC 10x Visium datasets with paired scRNA-seq datasets, we trained a k-NN classifier using the cell-type labels and coordinates of ST data and predicted cell-type labels for the localized coordinates of scRNA-seq data. We calculated ARI from the predicted and raw labels of the localized scRNA-seq data, which can measure localization accuracy globally.

#### Kullback-Leibler (KL) divergence

We adopted the KL divergence for the simulation experiments. We transferred the localized coordinates and raw coordinates of single cells to spatial grid density and calculated the KL divergence between them. KL divergence can measure the recovery performance of the spatial distribution density of cells.

#### Pearson correlation

We calculated the Pearson correlation of the localized and raw x-axis (and y-axis) coordinates of cells for the simulation experiments.

### Data availability

The data analyzed in this study can be accessed through: Human MTG scRNA-seq dataset: https://portal.brain-map.org/atlases-and-data/rnaseq/human-mtg-10x_sea-ad; Human M1 scRNA-seq dataset: https://portal.brain-map.org/atlases-and-data/rnaseq/human-m1-10x; Human DLPFC 10x Visium dataset: http://spatial.libd.org/spatialLIBD; Mouse brain cortex scRNA-seq dataset: https://www.dropbox.com/s/cuowvm4vrf65pvq/allen_cortex.rds; Mouse brain 10x Visium dataset: https://support.10xgenomics.com/spatial-geneexpression/datasets; Human SCC datasets: Gene Expression Omnibus (GEO) accession GSE144240; Mouse hippocampus scRNA-seq dataset: GEO accession GSE116470, or https://www.dropbox.com/s/cs6pii5my4p3ke3/mouse_hippocampus_reference.rds; Mouse hippocampus Slide-seqV2 dataset: Single Cell Portal (SCP) accession SCP815, Puck_200115_08; Mouse olfactory bulb scRNA-seq dataset: GEO accession GSE121891; Mouse olfactory bulb Slide-seqV2 dataset: SCP accession SCP815, Puck_200127_15; Mouse VISp scRNA-seq dataset: https://www.dropbox.com/sh/q687ajyvnkbtwkc/AACxlJAsJaJ5DVe-2mx3cbW2a?dl=0; Mouse brain cortex seqFISH+ dataset: https://github.com/CaiGroup/seqFISH-PLUS; Mouse visual cortex STARmap dataset: https://www.dropbox.com/sh/f7ebheru1lbz91s/AADm6D54GSEFXB1feRy6OSASa/visual_1020/20180505_BY3_1kgenes?dl=0&subfolder_nav_tracking=1.

### Code availability

The implementation of STALocator can be downloaded from https://github.com/zhanglabtools/STALocator.

## Acknowledgments

This work has been supported by the National Key Research and Development Program of China [No. 2021YFA1302500 to S.Z.], the National Natural Science Foundation of China [Nos. 32341013, 12326614, 12126605], the Key-Area Research and Development of Guangdong Province [No. 2020B1111190001], and the CAS Project for Young Scientists in Basic Research [No. YSBR-034 to S.Z.].

## Author contributions

Shihua Zhang conceived and supervised the project. Shang Li and Qunlun Shen designed and implemented the STALocator algorithm. Shang Li and Qunlun Shen validated the study. Shang Li, Qunlun Shen and Shihua Zhang wrote the manuscript. All authors read and approved the final manuscript.

