## Supplemental Figures for "Spatial Transcriptomics-Aided Localization for Single-Cell Transcriptomics with STALocator"

**Supplementary Figures**

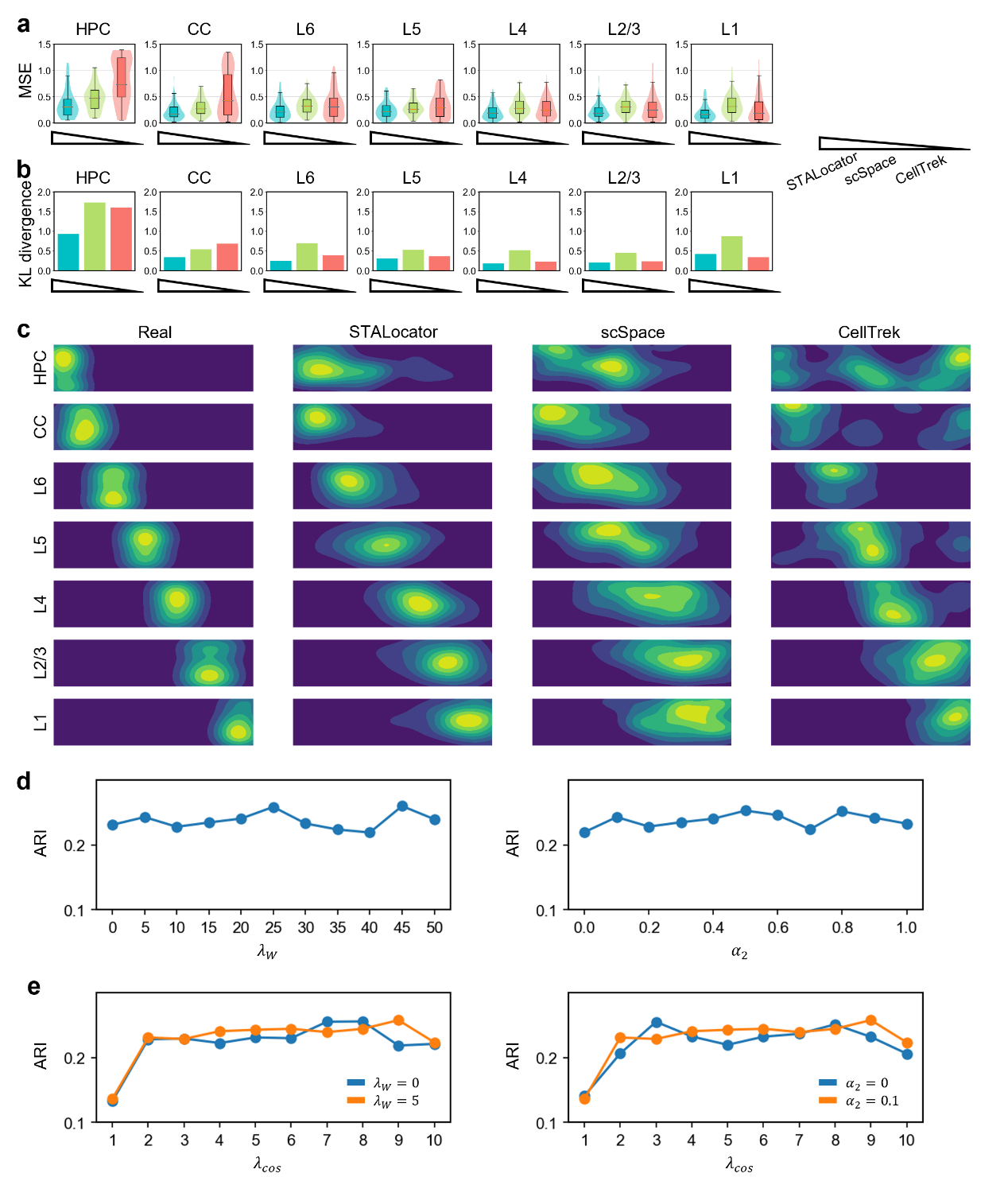

**Fig. S1. Evaluation of the recovery performance of each domain and analysis of roles of selected parameters. a,** Boxplot and density curve of localization error in terms of MSE for different methods. **b,** Bar chart of KL divergence calculated from spatial grid density for different methods. **c,** Contour plot showing spatial grid density for ground-truth and different methods. **d,** Sensitivity analysis of selected parameters. **e,** Ablation experiments of selected parameters.

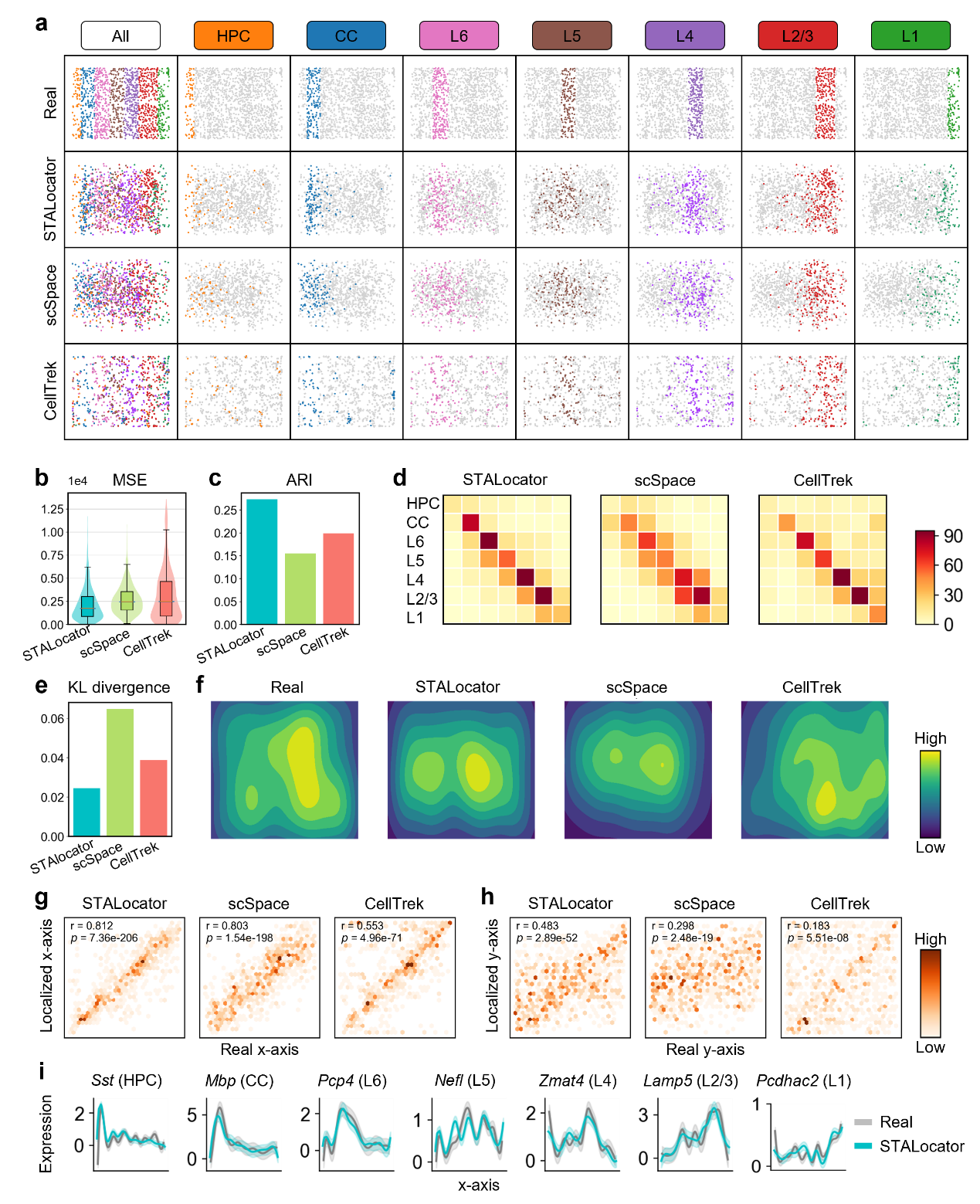

**Fig. S2. Evaluation of the recovery performance of the simulation experiment on the mouse visual cortex STARmap dataset with another resolution (700-pixel resolution). a,** Ground-truth annotation of seven domains and localization results for different methods. **b,** Boxplot and density curve of localization error in terms of MSE for different methods. **c,** Bar chart of localization accuracy measured by ARI for different methods. **d,** Confusion matrix showing correspondence between localized and raw labels for different methods. **e,** Bar chart of KL divergence calculated from spatial grid density for different methods. **f,** Contour plot showing spatial grid density for ground-truth and different methods. **g,** Density plot of the correlation test of the X-coordinates between different methods and ground truth. **h,** Density plot of the correlation test of the Y-coordinates between different methods and ground truth. **i,** The fitted expression trend along the x-axis of marker genes for STALocator and ground truth.

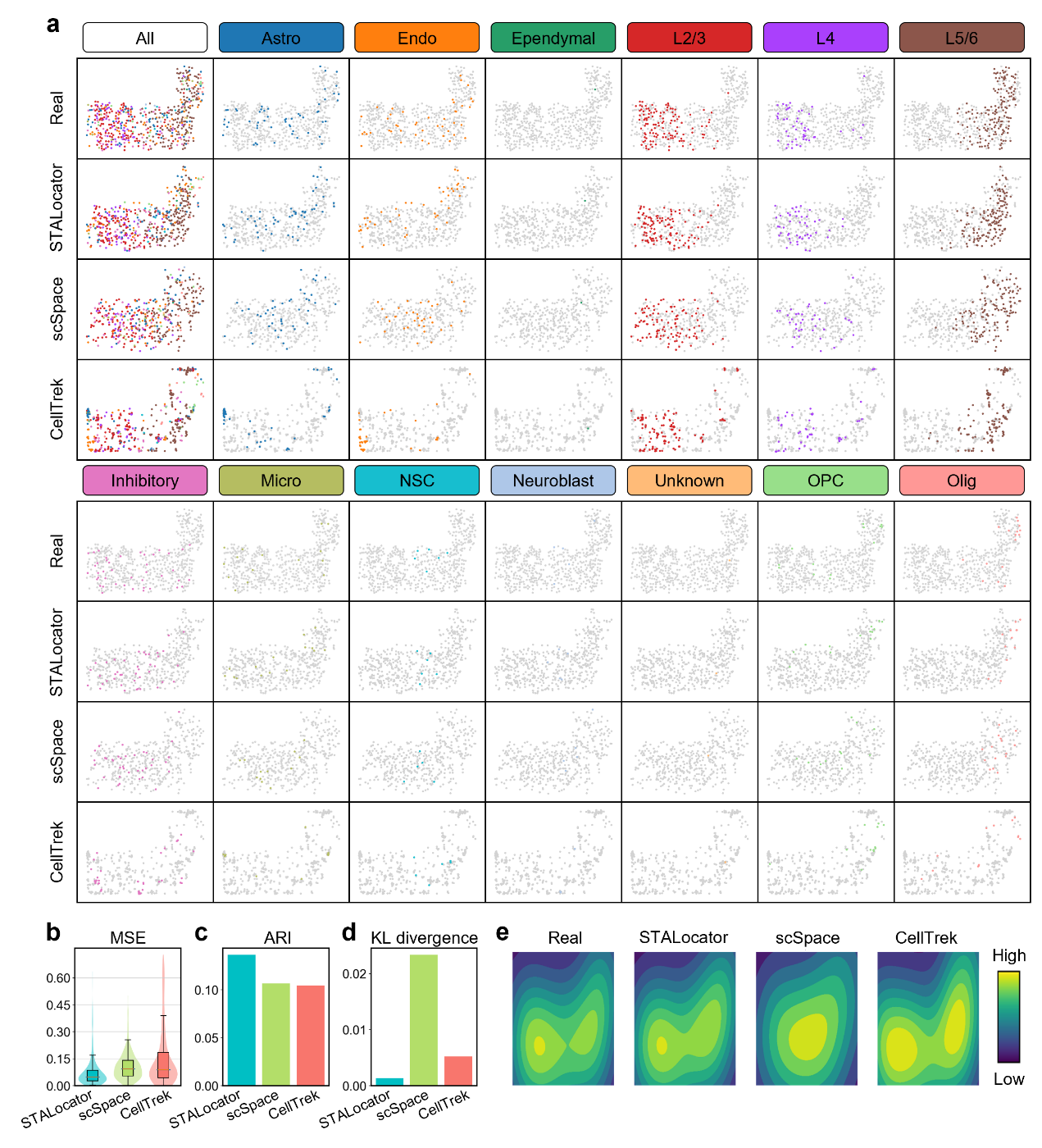

**Fig. S3. Evaluation of the recovery performance of the simulation experiment on the mouse brain cortex seqFISH+ dataset (500-pixel resolution). a,** Ground-truth annotation of seven domains and localization results for different methods. **b,** Boxplot and density curve of localization error in terms of MSE for different methods. **c,** Bar chart of localization accuracy measured by ARI for different methods. **d,** Bar chart of KL divergence calculated from spatial grid density for different methods. **e,** Contour plot showing spatial grid density for ground-truth and different methods.

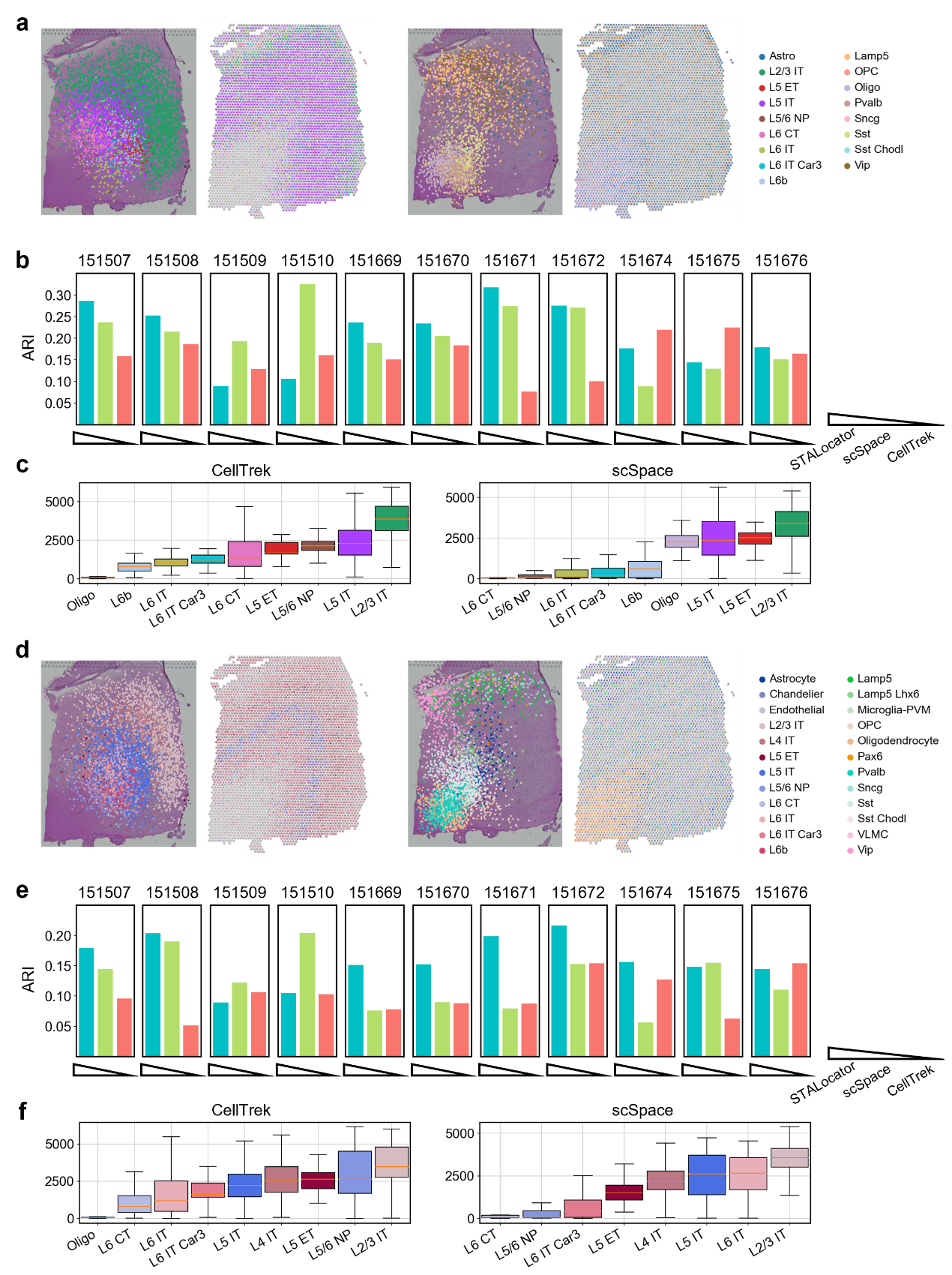

**Fig. S4. Comprehensive evaluation of the localization performance on the paired human brain cortex scRNA-seq datasets and human DLPFC ST dataset. a-c,** Localization performance evaluation on human brain M1 scRNA-seq dataset. **d-f,** Localization performance evaluation of human brain MTG scRNA-seq dataset. **a, d,** Spatial visualization of localization results of layer-associated cell types and other cell types. **b, e,** Bar chart of the recovery degree of the laminar organization based on ARI for all other sections. **c, f,** Boxplot of distance between WM and selected cell populations of localized scRNA-seq datasets by CellTrek and scSpace.

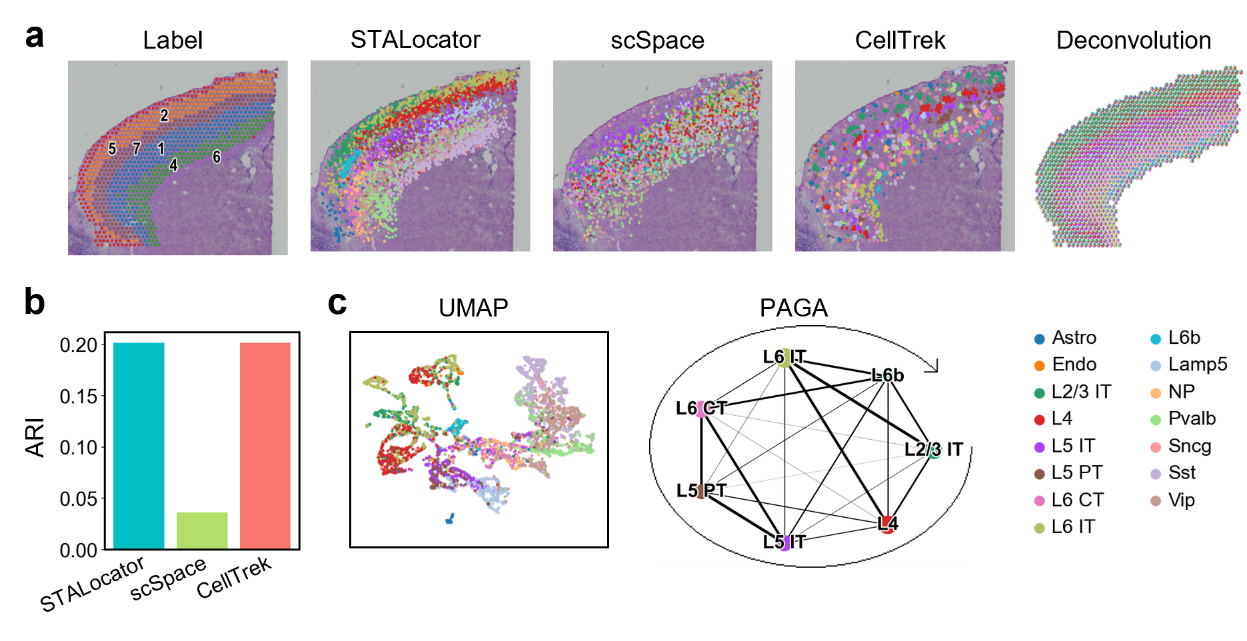

**Fig. S5. Evaluation of the localization performance on the paired mouse visual cortex scRNA-seq dataset and the mouse brain 10x Visium dataset.** **a,** Spatial visualization of tissue domain labels, localization results of different methods, and deconvolution results of RCTD. **b,** Bar chart of the recovery degree of the laminar organization based on ARI for different methods. **c,** UMAP visualizations and PAGA graphs generated from the STAGATE-derived low-dimensional representations of localized scRNA-seq dataset.

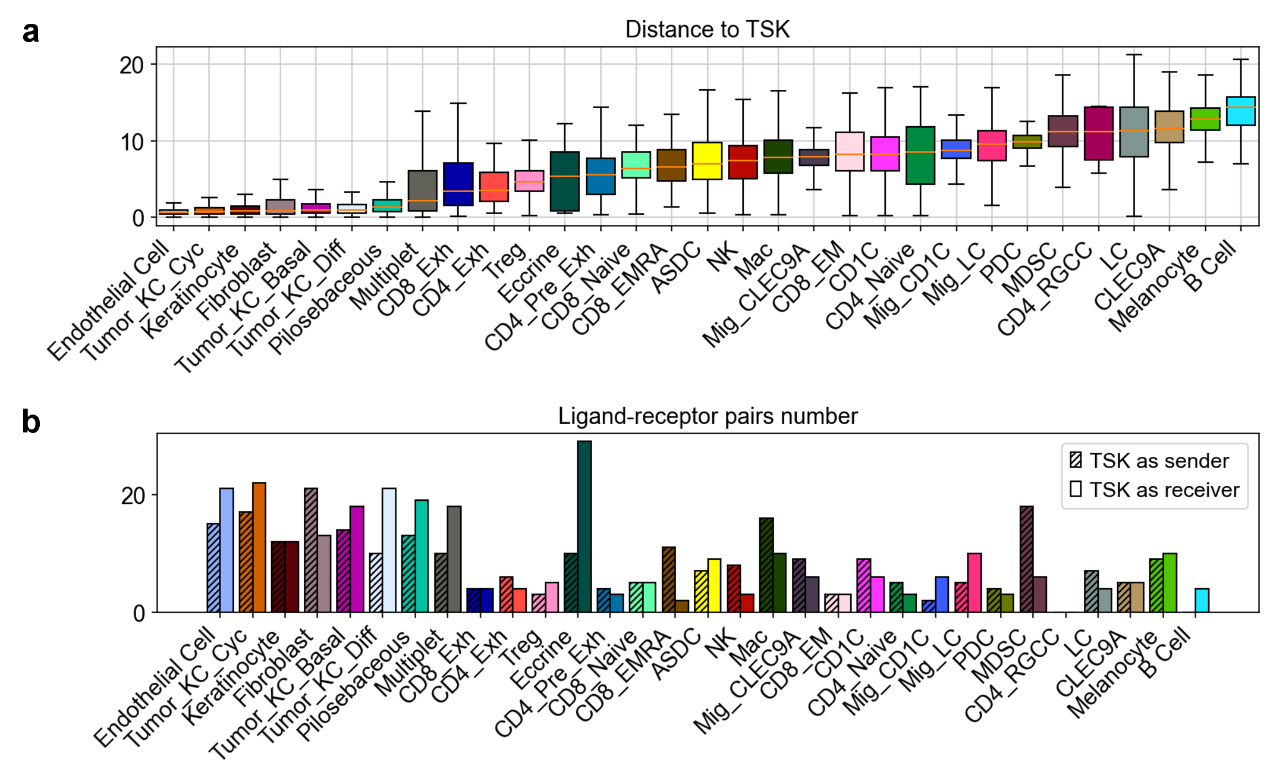

**Fig. S6. Analysis results of all cell subpopulations on the human SCC datasets. a,** Boxplot of the distance between TSKs and all cell subpopulations. **b,** Bar chart of ligand-receptor pairs numbers between TSK and all cell subpopulations.

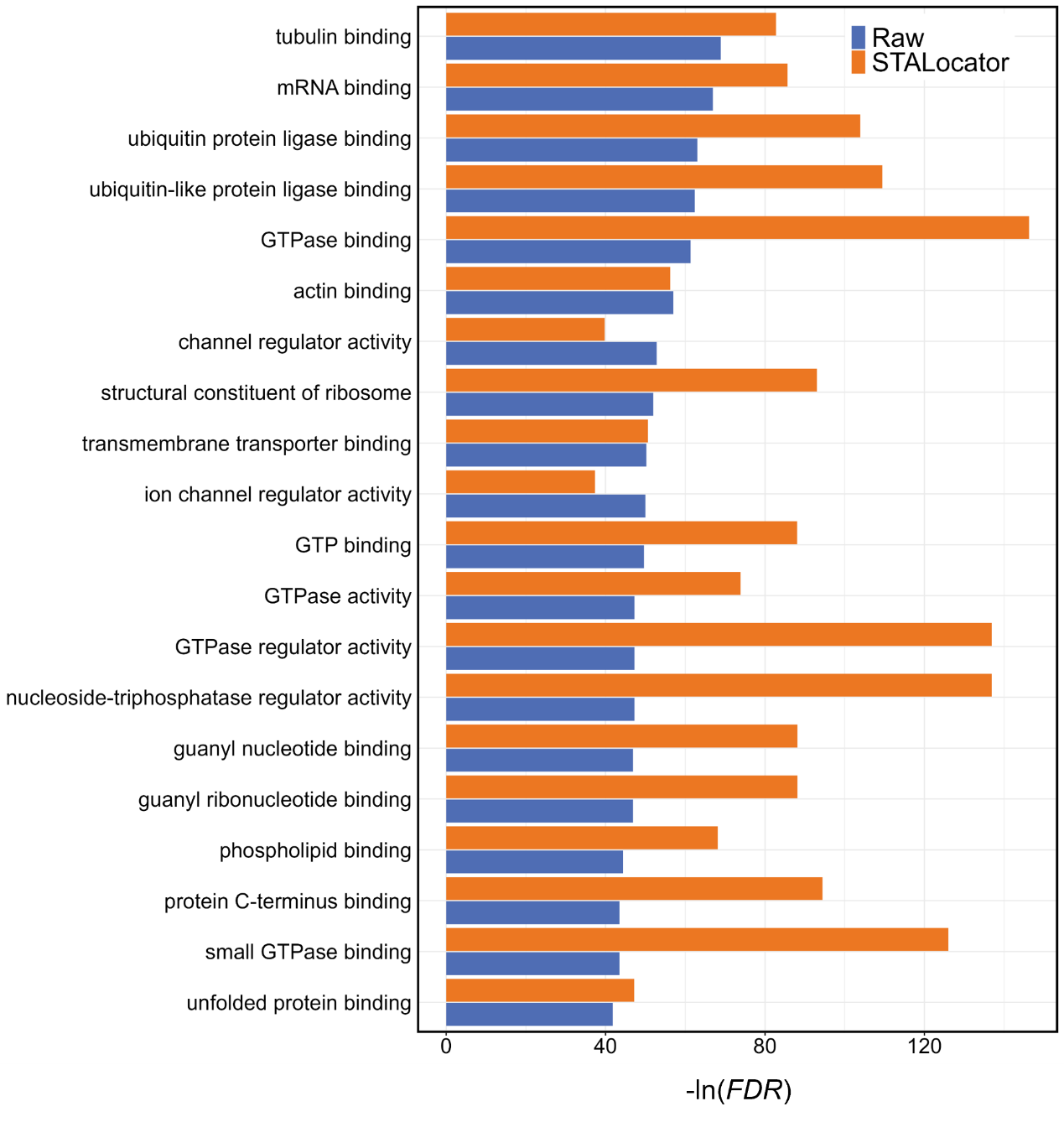

**Fig. S7. Shared GO terms of SVGs identified from the raw and enhanced mouse hippocampus Slide-seq dataset.**

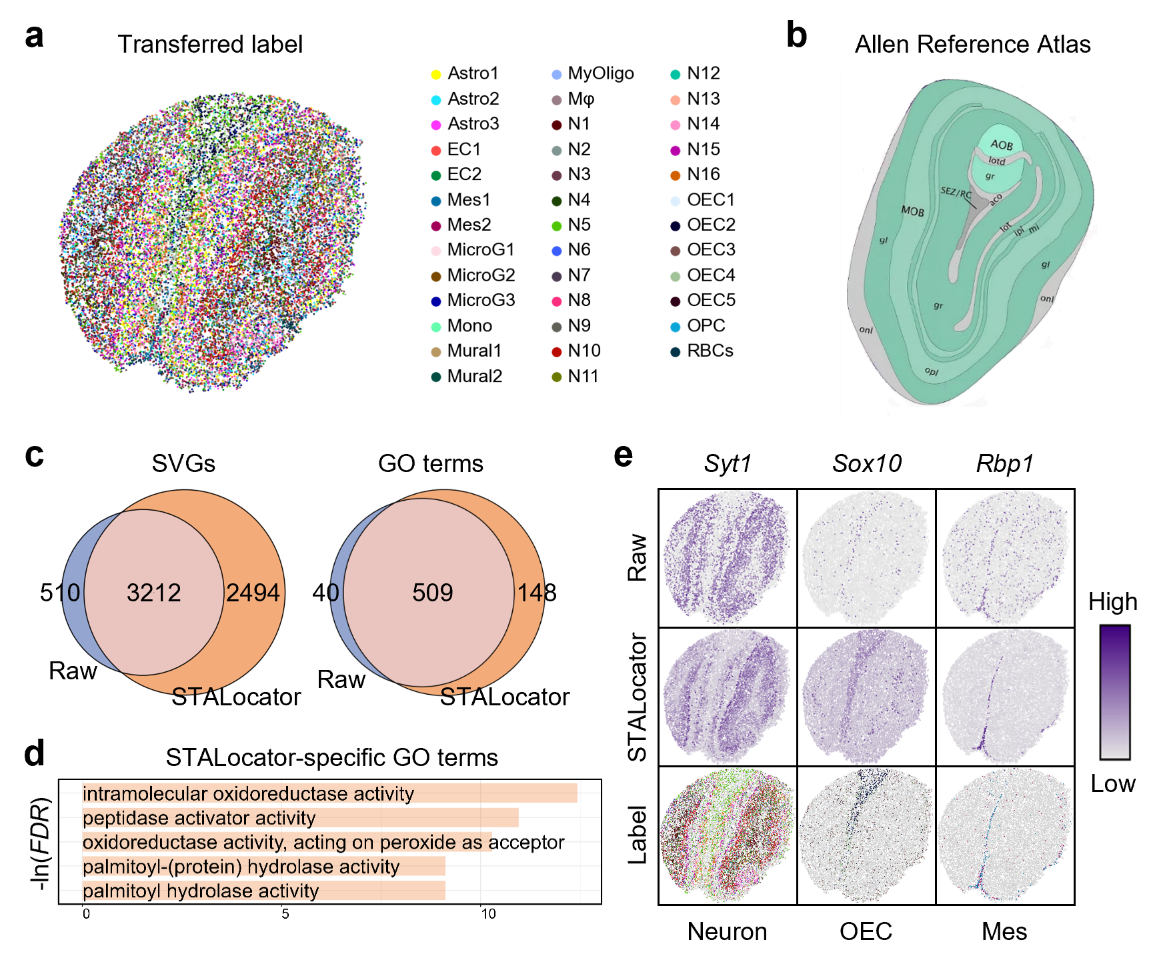

**Fig. S8. Application on the paired scRNA-seq dataset and Slide-seq dataset for mouse olfactory bulb. a,** Spatial visualization of cell-type labels transferred. **b,** ISH image from the Allen Brain Atlas. **c,** Venn plot of identified SVGs and enriched GO terms of the raw and enhanced dataset. **d,** Bar chart of the top five STALocator-specific GO terms. **e,** Spatial visualization of STALocator-specific SVGs of the raw and enhanced dataset and corresponding transferred labels.

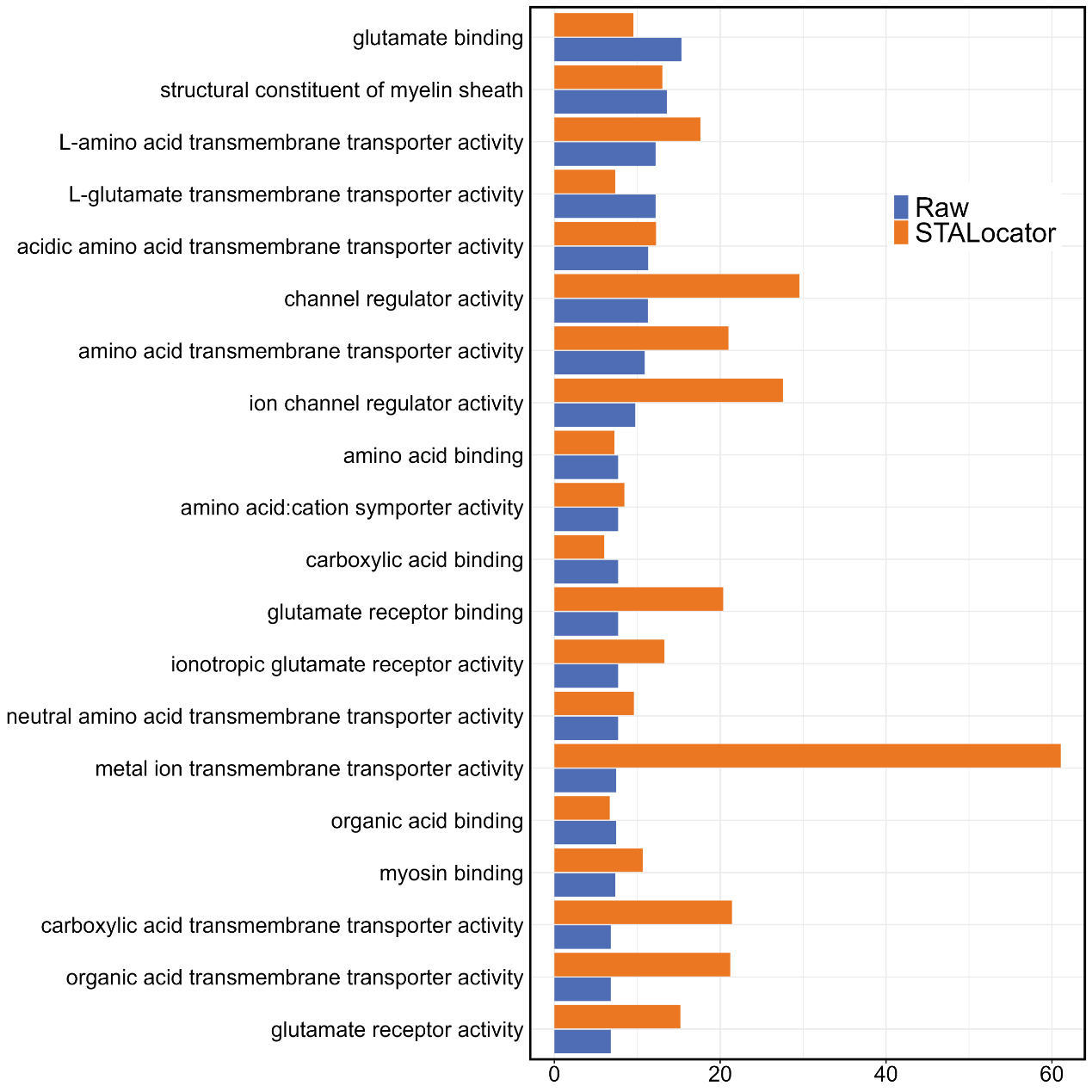

**Fig. S9. Shared GO terms of SVGs identified from the raw and enhanced mouse visual cortex STARmap dataset.**

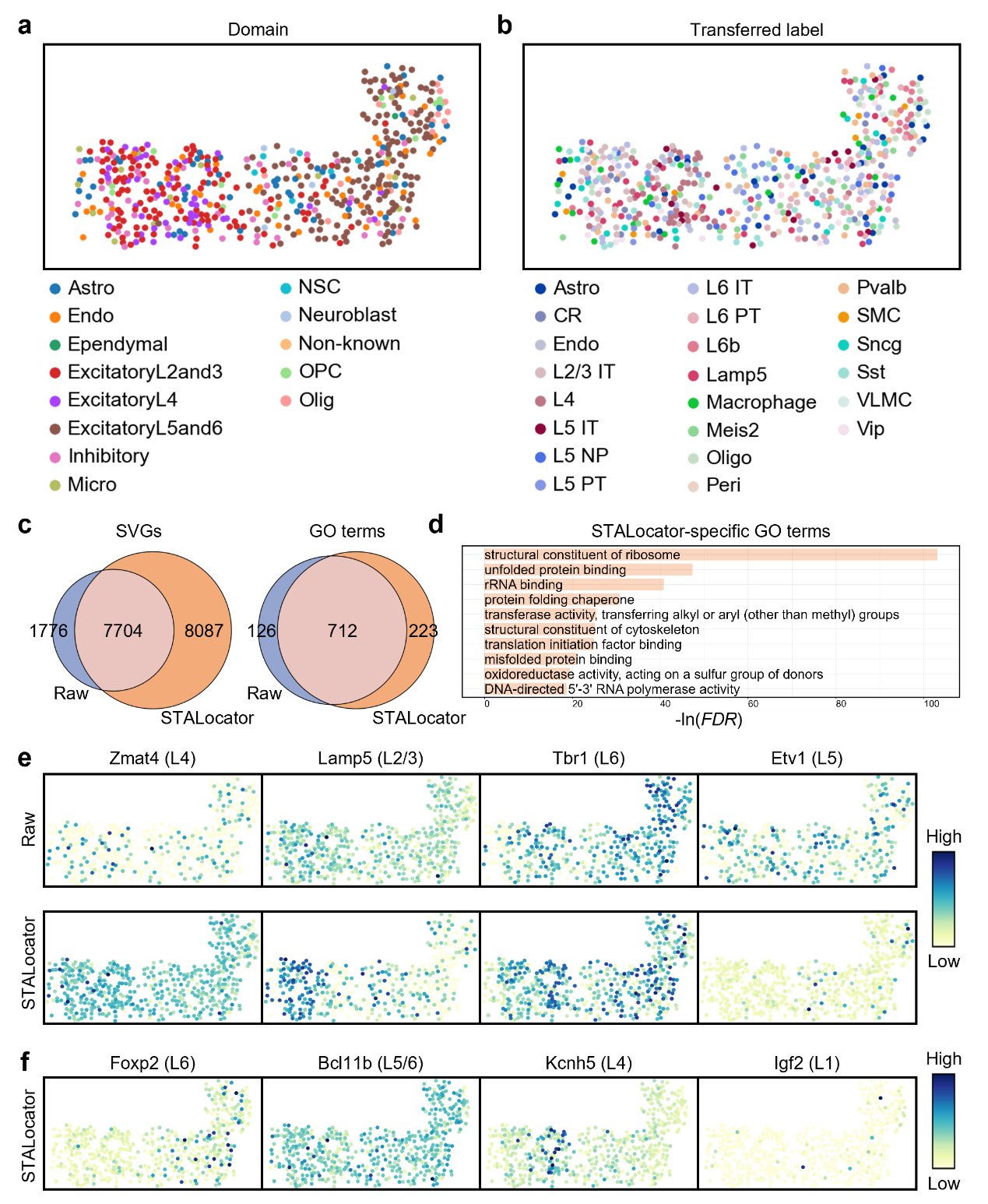

**Fig. S10. Application on the paired mouse VIsp scRNA-seq dataset and the mouse brain cortex seqFISH+ dataset. a,** Spatial visualization of manual annotation. **b,** Spatial visualization of cell type labels transferring results. **c,** Venn plot of identified SVGs and enriched GO terms of the raw and enhanced dataset. **d,** Bar chart showing the top ten STALocator-specific GO terms. **e,** Raw and recovered expression profiles of measured marker genes. **f,** Predicted expression of unmeasured marker genes.

**Supplementary Tables**

**Table S1. Description of all scRNA-seq datasets used in this study.**

| **Platform/Protocol** | **Tissue** | **#Cells** | **Related figures** | **Reference** |
| --- | --- | --- | --- | --- |
| **STARmap** | Mouse visual cortex | 1207 | Fig 2,  Fig S1, S2 | ^1^ |
| **seqFISH+** | Mouse brain cortex | 523 | Fig S3 | ^2^ |
| **10x Chromium** | Human squamous cell carcinoma (SCC) | 26299 | Fig. 4,  Fig. S6 | ^3^ |
| **10x Chromium** | Human middle temporal gyrus (MTG) | 10000 | Fig. 3,  Fig. S4 | ^4^ |
|  | Human primary motor cortex (M1) | 10000 |  |  |
| **SMART-Seq2** | Mouse visual cortex | 4785 | Fig. S5 | ^5^ |
| **Drop-seq** | Mouse hippocampus | 20000 | Fig. 5,  Fig. S7 | ^6^ |
| **10x Chromium** | Mouse olfactory bulb | 20000 | Fig. S8 | ^7^ |
| **SMART-Seq2** | Mouse primary visual area (VISp) | 11759 | Fig. 6,  Fig. S9, S10 | ^8^ |

**Table S2. Description of all ST datasets used in this study.**

| **Platform** | **Tissue** | **Section** | **#Spots/Beads/Cells** | **Related figures** | **Reference** |
| --- | --- | --- | --- | --- | --- |
| **STARmap (binned)** | Mouse visual cortex | Res_900,  Res_700 | 127,  198 | Fig 2,  Fig S1, S2 | ^1^ |
| **seqFISH+ (binned)** | Mouse brain cortex | Res_500 | 71 | Fig S3 | ^2^ |
| **Spatial Transcriptomics** | Squamous cell carcinoma (SCC) | Patient 2 | 666 | Fig. 4,  Fig. S6 | ^3^ |
| **10x Visium** | human dorsolateral prefrontal cortex (DLPFC) | 151507,  151508,  151509,  151510,  151669,  151670,  151671,  151672,  151673,  151674,  151675,  151676. | 4226,  4384,  4789,  4634,  3661,  3498,  4110,  4015,  3639,  3673,  3592,  3460. | Fig. 3,  Fig. S4 | ^9^ |
|  | MouseBrain | Section 1  (Sagittal-Anterior) | 1074 | Fig. S5 | 10x Visium demo |
| **Slide-seqV2** | Mouse hippocampus | Puck_190921_21 | 10000 | Fig. 5,  Fig. S7 | ^10^ |
|  | Mouse olfactory bulb | Puck_200127_15 | 20139 | Fig. S8 |  |
| **STARmap** | Mouse visual cortex |  | 1207 | Fig. 6,  Fig. S9 | ^1^ |
| **seqFISH+** | Mouse brain cortex |  | 523 | Fig. S10 | ^2^ |
